# Electrophysiological features of signals recorded from white matter

**DOI:** 10.64898/2026.07.11.737939

**Authors:** Ryan Jafri, Fernando A. Ortega, Pranathi Manivannan, Zahra Jourahmad, Davin Devara, Layth S. Mattar, Sanjay Kirshna, Geoffrey Liu, Saipravallika Chamarti, Alica Goldman, Lu Lin, Vaishnav Krishnan, Atul Maheshwari, Garrett P. Banks, Mohammed Hasen, Danika L. Paulo, Andrew J. Watrous, Benjamin Y. Hayden, Jeffrey M. Yau, Sameer A. Sheth, Nicole R. Provenza, Nicholas Murphy, Sarah R. Heilbronner, Eleonora Bartoli

## Abstract

Intracranial neurophysiology studies have typically ignored signals from electrodes located in white matter (WM), assuming that their information content is artifactual or related to nearby gray matter (GM). Here, we tested the electrophysiological and functional features of signals recorded from different WM locations. Signals were recorded from 19 patients undergoing intracranial monitoring for drug-resistant epilepsy by means of stereo-electroencephalography (sEEG). Each sEEG electrode was classified into WM or GM based on the surrounding tissue. We obtained recordings from a total of 1,717 sEEG electrode contacts, 36% in WM, while the patients were in awake resting state (5 minutes). For each sEEG electrode, we employed a model-based spectral decomposition to separate periodic and aperiodic components, and we computed signal complexity metrics. For a subset of participants, we computed WM structural information from diffusion-weighted magnetic resonance imaging and we evaluated functional signals during a cognitive control task. Our results show that signals recorded from WM have different spectral features and higher complexity than GM. Complexity correlates positively with fractional anisotropy, and modulations related to behavior during the task were detected in WM. Overall, this indicates that WM signals carry information that may reflect signal propagation across WM fiber tracts.

## Introduction

Classically, neuroscientists studying functional signals in the brain have focused on gray matter (GM) — the collection of somas and axon hillocks that are responsible for generating action potentials. The signals that can be obtained from white matter (WM) — the myelinated axons that transmit information across short and long distances — have been largely ignored, and WM has been primarily considered for its structure. Evidence that functional information exists in WM has primarily come from functional magnetic resonance imaging (fMRI) studies, demonstrating that WM signals can be task-related, time-locked, and spatially modular, just like GM signals (Peer *et al*., 2017; Schilling *et al*., 2023; Spencer *et al*., 2025). Studies using intracranial neurophysiology, on the other hand, have typically masked out or otherwise ignored WM contacts, presuming that their information content is artifactual or a simple reflection of nearby GM. However, a few notable exceptions exist. One study using intracranial human recordings demonstrated that WM signals cannot be fully explained by nearby GM signals (Mercier *et al*., 2017), leading the authors to speculate that WM signals might be partially capturing information traveling on WM fiber tracts. Another study exploited the presence of differences in spectral content between GM and WM signals to perform semi-automated classification of electrodes positions (Greene *et al*., 2021), although the primary objective of the study was to provide means to discard WM signals. Recent work found that functional connectivity between WM signals measured intracranially and by fMRI was strongly correlated at a single-participant level (Huang *et al*., 2023). The inclusion of recordings from WM locations, in addition to GM ones, improved movement decoding performance, with clear implications for brain-computer interface applications (Li *et al*., 2021). Crucially, WM signals were recently shown to carry information related to seizure propagation across seizure networks in patients with epilepsy (Revell *et al*., 2026), demonstrating the presence of clinically-relevant signals in WM and reviving the interest in WM signal features (Mercier, 2026).

Together, this evidence suggests that WM signals may be influenced by a mixture of sources. These sources may include signals from nearby GM and signals unique to WM, potentially arising from activity occurring along WM tracts, offering clinically relevant insights into local and interconnected circuits. Here, we tested whether intracranial signals recorded from WM and GM have other distinct spectral features and information content, and if this relates to specific WM fiber tracts. We hypothesized that WM signals carry network-wide information content reflecting the integration of activity from interconnected GM areas. Although some locations in the WM are dominated by axons from nearby GM, others contain long-range passing fibers that transmit information from distant brain regions. In other words, WM signals are not simply attenuated local field potentials from nearby GM. To test this hypothesis, we employed intracranial recordings in patients undergoing invasive monitoring for pharmaco-resistant epilepsy by means of sEEG. We determined whether WM signals differ from GM ones in more ways than expected by a simple signal attenuation (Greene *et al*., 2021) by evaluating spectral features, signal complexity metrics, and WM signals’ association with structural information from diffusion-weighted magnetic resonance imaging (dMRI). Finally, we evaluate whether functional signals are present in WM signals by testing for behavior-related modulations during a stop-signal task.

## Materials and Methods

### Human Subjects

Nineteen subjects (10 females, age ranging from 20 – 63 years, mean and standard deviation: 38 ± 13 years) with drug-resistant epilepsy consented to participate in research while they underwent invasive intracranial activity monitoring with sEEG at Baylor St. Luke’s Medical Center (Houston, Texas, USA). Experimental procedures were conducted in accordance with the policies and principles outlined in the Declaration of Helsinki and were approved by the Institutional Review Board at Baylor College of Medicine (H-18112). Patients provided written and verbal consent.

### Stereo-EEG Electrodes

Patients were implanted with a variable number of sEEG probes based on clinical needs (median number of probes = 15). The sEEG probes had either a 0.8 mm diameter, with 8–16 electrode contacts of 2 mm length and a 3.5 mm center-to-center distance (PMT Corporation, MN, USA), or a 1.28 mm diameter, with nine electrode contacts of 1.57 mm length and a 5.0 mm center-to-center distance between contacts (AdTech Medical Instrument Corporation, WI, USA).

### Electrode localization and assignment to GM or WM

We determined electrode locations for each subject by using the software pipeline intracranial Electrode Visualization, iELVis (Groppe *et al*., 2017). In short, the postoperative CT image was registered to the preoperative T1 anatomical MRI image using FSL (Jenkinson *et al*., 2012). The location of each electrode contact was then identified in the CT-MRI overlay using BioImage Suite (Papademetris *et al*., 2006). The coordinates for each electrode contact were used to identify their position with respect to anatomical boundaries by mapping the Destrieux Cortical Atlas to the MRI of each patient (Destrieux *et al*., 2010). Labels from voxels within 3 mm of each coordinate were used to classify each electrode as GM, WM, subcortical, or boundary (if there was a mix of GM and WM voxels within the radius; Figure 1A). For analyses, only signals from electrode contacts classified as GM and WM were retained (Figure 1B). For group-level visualization purposes (Figure 1C), we used the MNI305 coordinates and the *fsaverage* cortical surface from Freesurfer (version 7.4.1)(Dale, Fischl and Sereno, 1999).

**Figure 1.**
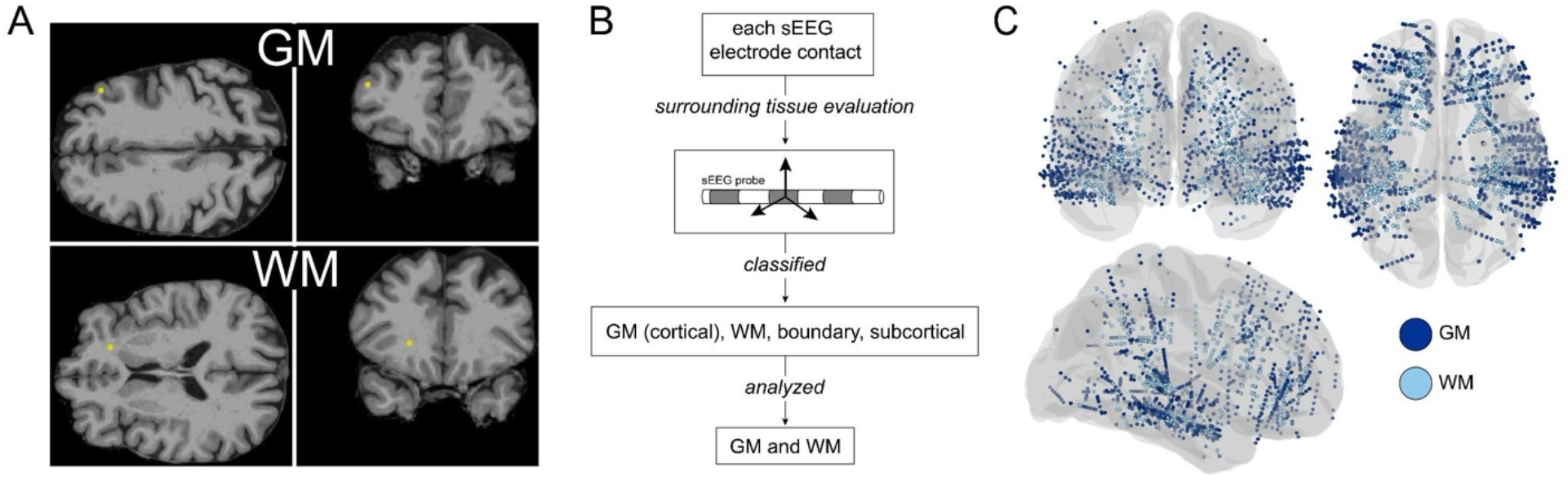
Electrode assignment to WM and GM. Panel A: example of two sEEG electrode contact locations (same patient, same sEEG probe), one in cortical GM and one in WM; the electrode contacts are represented by a yellow circle. Panel B: schematic of the assignment to GM, WM, boundary or subcortical based on the tissue surrounding each electrode contact, using a 3 mm radius, represented by the arrows. Panel C: WM and GM electrode locations used for analysis for all participants, displayed on the fsaverage brain.

### Electrophysiological Recordings

Awake resting-state sEEG signals were recorded by a 256-channel BlackRock Cerebus system (BlackRock Microsystems) with 2 kHz sampling rate and a fourth-order Butterworth bandpass filter of 0.3–500 Hz. Two contacts visually determined to be in WM were used for ground and reference. The recordings were obtained while the patients were at the epilepsy monitoring unit, at least two days after surgery and consisted of 5-minute resting-state recordings (eyes-open, fixation cross displayed on a monitor). Signals were processed in MATLAB (R2019a, MathWorks, MA, USA) and they were inspected for line noise, recording artifacts (e.g., contamination by muscle activity, presence of large deflections, additional noise components), and interictal epileptic activity. Electrodes with noise, artifacts and epileptic activity were excluded from further analysis. Next, the signal from each electrode was re-referenced to the average of all electrodes that survived exclusion criteria and down-sampled to 1kHz. A full set of control analyses with an alternative re-referencing scheme is reported in the related result subsection. In total, we obtained signals from 1,717 electrode contacts for the remaining analyses (64% from GM, 36% from WM). See Supplementary Information for electrode contacts numbers for each participant, Figure 1C for their position with respect to a template brain.

### Power Spectrum Modeling

For each signal, we computed the power spectrum using the Welch method (3-second sliding windows, 50% overlap) for each contact. We used an established approach (*specparam*, formerly *fooof*) (Donoghue *et al*., 2020) to model the power spectrum (0.3-250 Hz) into a combination of aperiodic components, capturing the decay of power with increasing frequencies (1/f shape) and periodic components, employing custom frequency ranges. The aperiodic components included offset (y-intercept), knee (bend in the 1/f profile), and exponent (parameter *x* in the 1/f^*x*^ model fit, capturing the rate of decay of the 1/f profile). To estimate periodic components, we divided the frequency ranges in contiguous, non-overlapping bands: delta (0.3-4 Hz), theta (4-8 Hz), alpha (8-13 Hz), beta (13-30 Hz) using the modified algorithm Spectral Parameterization for the Broadband Analysis of Neural Data (SP-BAND) (Phillips, Teimouri and Aazhang, 2024). This algorithm has the advantage of constraining the estimates of periodic components into canonical bands, which simplified our band-specific comparison approach for WM and GM estimates. In the model, we also included two high-frequency ranges 30-50 and 50-250 Hz to improve the overall goodness of fit of the model given the presence of peaks due to line noise and its harmonics that would not be otherwise captured. Given that there were no physiological oscillations in this range, these ranges were included just to improve model fit and were not further analyzed. Periodic components were fit in natural log space; each frequency band was split into 1 division; prominence value set to 0.5; line noise cancellation at 60 Hz, and peak fitting process was not iterated.

### Complexity Measures

For each contact, we computed two measures of signal complexity: Higuchi Fractal Dimension (HFD) and Lempel-Ziv Complexity (LZC). Signal complexity can be artificially inflated by the presence of noise in the data. We reduced the potential influence of artifacts related to motion or electrical noise by identifying a stable two-minute portion of the recording for each patient. To standardize the vertical range of the signal between subjects the signal was z-scored within each subject (Murphy *et al*., 2023).

HFD is a measure of self-similarity that allows us to estimate the variety of neural systems contributing to a given signal, where a greater value indicates a stronger departure from homogeneous neural activity (Shamsi, Ahmadi-Pajouh and Seifi Ala, 2021; Grosu *et al*., 2023). We estimated HFD using N = 8 iterations (kmax parameter).

The LZC complements the HFD and other measures of non-linear activity by providing a view of the total information content without focusing extensively on how the information is structured (Ibáñez-Molina *et al*., 2015). The LZC estimates the number of unique patterns in the binarized time-series and is summarized by the number of iterations required to complete the estimation. Samples exceeding the mean of the rectified time series were coded as “1”, those less than the mean were coded as “0”. Normalized log complexity was estimated using an implementation (*calc_lz_complexity* function by Quang Thai, on MATLAB Central File Exchange, 2026) of the LZC algorithm (Lempel and Ziv, 1976).

### DMRI Acquisition and preprocessing

Prior to surgical implantation of the sEEG electrodes, a subset of patients (n=9) underwent anatomical imaging at the Core for Advanced MRI at Baylor College of Medicine. Diffusion-weighted imaging data was acquired on a Siemens Prisma 3T system for 2 b-values (b = 1000, 2000 s/mm2) with two phase-encoding directions (anterior-to-posterior and posterior-to-anterior) using a dual-echo echo-planar protocol (Repetition time = 3400 ms, Echo time = 85.8 ms, Voxel size: 1.5 mm isotropic, Multiband acceleration factor = 4, 99 slices). A total of 7 interleaved non-diffusion-weighted (b0) volumes were collected, and 92 diffusion-sensitizing gradient directions were applied per acquisition.

DMRI data were preprocessed using FSL (v6.0.7.10; FMRIB, Oxford, UK). Susceptibility-induced off resonance field estimation and correction were performed using TOPUP, which estimated the distortion field from a pair of b0 volumes acquired with opposing phase-encoding directions (AP and PA). The estimated field was subsequently used to unwarp the full AP dMRI series using EDDY, which simultaneously corrected for eddy current-induced distortions and subject motion. The b = 1000 s/mm^2 shell was then extracted from the eddy corrected data using FSL’s *select_dwi_vols*, and the diffusion tensor was fit voxelwise using DTIFIT. Maps of fractional anisotropy (FA) were derived from the resulting eigenvalue and eigenvector decomposition.

Registration of the pre-operative T1-weighted structural image to subject-specific dMRI space was performed using FLIRT (FSL Linear Image Registration Tool). To maximize cross-modality registration accuracy, the transform was estimated between the brain-extracted T1 and the mean b0 volume, as it provides substantially greater soft-tissue contrast for rigid-body alignment across modalities. Prior to registration, both the T1 and b0 images were reoriented to standard orientation using *fslreorient2std*. A 6-degree-of-freedom (rigid-body) transformation was estimated using normalized mutual information as the cost function. The resulting transformation matrix was subsequently applied to additionally resample the T1 into the FA image grid for visualization and quality control.

### sEEG Electrode ROI Construction and Mapping

Pre-implantation sEEG electrode coordinates were provided in FreeSurfer tkRegister RAS space, defined relative to the pre-operative T1 volume. Single voxel binary regions of interest (ROIs) were generated for each white matter electrode contact by converting the tkRegister RAS coordinates to voxel indices in T1 space using the FreeSurfer vox2ras-tkr convention, implemented from the geometry of the T1 NIfTI header. Each ROI was then mapped from T1 space into native diffusion (FA) space using *img2imgcoord* (FSL), which applies the previously estimated FLIRT transformation matrix at the level of individual voxel coordinates. ROIs were reoriented to standard orientation prior to mapping to ensure consistency with the reoriented reference images used during registration. Each T1-space voxel coordinate was transformed and rounded to the nearest voxel in FA space. Each electrode ROI was subsequently dilated to a spherical mask of 2 mm radius. The structuring element was constructed in millimeter space using the image voxel dimensions, such that the resulting sphere is geometrically accurate regardless of voxel anisotropy. Dilation was performed using binary morphological dilation (*scipy*.*ndimage*; Python). Mean FA was then extracted from each spherical mask. Analysis scripts used in this pipeline are publicly available at https://github.com/Thenandolab/sEEG-Diffusion-ROI-Pipeline.

### Stop-Signal Task Analyses

For a subset of patients (n=9), we further analyzed signals from WM locations during the performance of a stop-signal task, adapted from the “STOP-IT2_beta03” software (Verbruggen, Logan and Stevens, 2008). Briefly, the stop-signal task requires participants to respond quickly to the presentation of a visual go cue (right or left arrow) using a response pad, and to withhold the response when the go cue is followed by a stop cue (change in color of the arrow, turning yellow occurring on 25% of trials, here leading to 72 stop trials). The delay between go and stop cues was varied adaptively using standard procedures to identify the delay leading to the ability to stop the response in 50% of the trials, thus obtaining a comparable number of Stop-Success and Stop-Fail trials. Extended details about the paradigm can be found in our previous publication (Kang *et al*., 2025). Signals from WM locations were down-sampled to 1 kHz and spectrally decomposed using a family of Morlet wavelets (7 cycles), with center frequencies ranging from 2 to 150 Hz in 1 Hz steps. We identified the time of the stop signal presentation (aligned to the neural data through a photoresistor) and we selected time-frequency power values from 500 ms before to 600 ms after the stop cue. Power values during each stop trial were normalized to percent change with respect to a baseline period (between 1000 to 500 ms before stop cue presentation). Power % change values were computed for the WM locations employed in the resting state analysis, after excluding electrode locations displaying noise or artifacts.

### Statistical Analyses

All resting-state derived measures were modeled with a mixed-effects model approach (*lmer* function in R) with fixed effects for matter type (levels: GM and WM) and random effects for participants, and electrode contacts nested within participants. Correlations between metrics were determined by computing Spearman’s rank correlation coefficient. To assess the presence of differences in the time-frequency power values for the Stop-signal task, we employed permutation testing (5,000 permutations) randomly shuffling labels between Stop-Success and Stop-Fail trials. This approach allowed us to estimate the distribution of differences at each time-frequency point under the null hypothesis. To account for multiple comparisons across the time-frequency space (1100 by 150 matrix of time-frequency values) we employed a cluster-based correction for multiple comparisons at a threshold of 0.05. For all analyses, exact p-values are reported for values larger than 0.001. Statistical significance is denoted as *p<0.05, **p<0.01, and ***p<0.001 in the figures.

## Results

### Aperiodic and periodic activity differences between WM and GM

Spectral features obtained from the parametrization of the power spectra revealed that signals recorded from WM locations had lower offset values than GM (mean and standard deviation for WM offset = 2.97 ± 0.902; GM offset = 3.88 ± 0.861; β = −0.91, p<0.001) and a shallower 1/f shape (WM exponent = 2.46 ± 0.388; GM exponent = 2.77 ± 0.374; β = −0.3, p<0.001; Figure 2A). Overall, these results indicate that WM signals had lower overall power but also a different distribution of power across frequencies, with a higher concentration of power at higher frequencies versus low frequencies in WM signals. The exponent was lower for WM versus GM (i.e., shallower 1/f shape for WM) consistently for all participants (Figure 2B).

**Figure 2.**
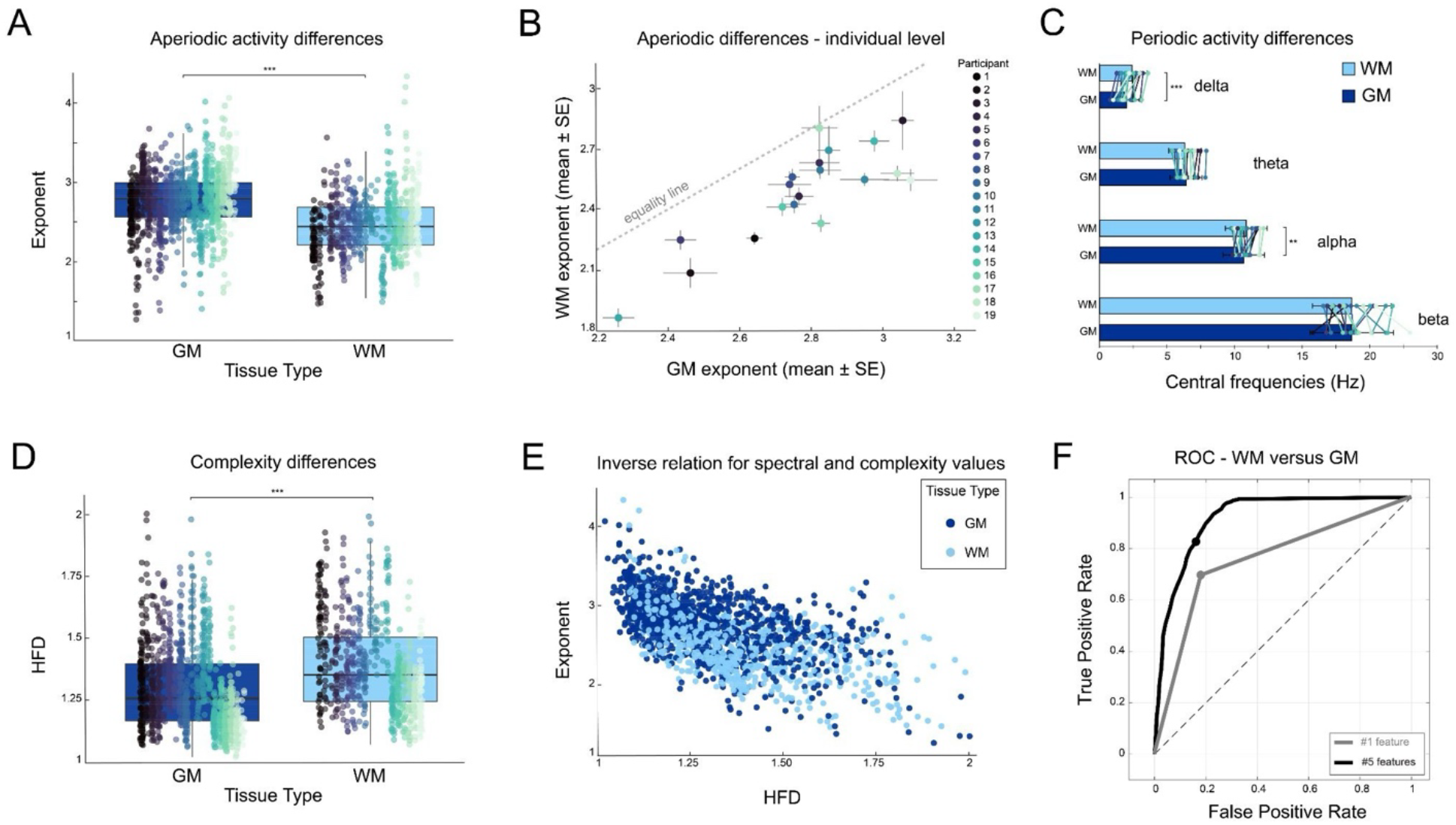
Spectral and complexity differences between WM and GM. Panel A: aperiodic activity values (exponent) are displayed as a boxplot for WM and GM, with the value for each electrode overlaid as a circle, ordered and color-coded by the participant number. B: exponent values for GM (x-axis) and WM (y-axis) for each participant are averaged across all electrodes for the participant (circle) and displayed with their standard error (SE, vertical and horizontal bars around circle). The dashed line denotes the equality line, with all values falling below it, denoting that the WM exponents are lower than GM for all participants. Panel C: central frequency values (x-axis) for canonical rhythms (delta, theta, alpha, beta) are displayed for WM and GM, with each bar showing the overall average and standard deviation. The average values for each participant are displayed as overlaid points connected by lines, color coded as in panel B. Panel D: HFD is displayed as a boxplot for WM and GM, with the value for each electrode overlaid as a circle, ordered and color-coded by the participant number as in panel B. Panel E: inverse relationship between HFD (x-axis) and exponent values (y-axis) for each electrode (circles, color coded by WM and GM assignment). Panel F: ROC curves for the classifier using one feature (AUC = 0.76) and 5 features (AUC = 0.92), showcasing the ability to classify electrodes into WM and GM. For all panels, asterisks denote significance derived from the mixed-effects model contrast between GM and WM, with *p<0.05, **p<0.01, and ***p<0.001.

When considering periodic features (Figure 2C), low frequency signals recorded from WM had a significantly higher central frequency, corresponding to faster oscillations in the delta range (WM delta range central frequency = 2.40 ± 1.13 Hz; GM delta range central frequency = 2.00 ± 1.12 Hz; β = 0.43, p<0.001). The effect was also present, to a smaller extent, in the alpha range (WM alpha range central frequency = 10.9 ± 1.55 Hz; GM alpha range central frequency = 10.7 ± 1.54 Hz; β = 0.21, p=0.005). No other reliable differences in central frequency estimates were detected for the other ranges considered (WM theta range central frequency = 6.32 ± 1.20 Hz, GM = 6.41 ± 1.19 Hz, β = 0.03, p=0.63; WM beta range central frequency = 18.7 ± 2.92, GM = 18.7 ± 3.10, β = −0.18, p=0.21).

### Higher complexity for WM signals

Complexity was higher for signals recorded from WM locations, both when considering HFD (HFD WM: 1.39 ± 0.187; HFD GM: 1.30 ± 0.171; β = 0.107, p<0.001; Figure 2D) and LZC (LZC WM: 0.464 ± 0.277; LZC GM: 0.354 ± 0.239; β = 0.120, p<0.001). Furthermore, complexity inversely correlated with the exponent of the power spectra, such that higher complexity scores were associated with lower exponent values, corresponding to a shallower slope (HFD-exponent rho = −0.59, p<0.001; LZC-exponent: rho = −0.18, p<0.001; Figure 2E). To test if the inverse correlation was robust at an individual level, we repeated the analysis for each participant separately. The negative correlation between HFD and exponent was significant for each participant (HFD-exponent rho ranging from: −0.28 with p=0.03, to: −0.94, with p<0.001). For LZC-exponent, the correlation was significant for 15 out of 19 subjects (LZC-exponent rho ranging from: −0.10, with p=0.43, to −0.73, with p<0.001).

### Decoding of WM & GM signals based on spectral and complexity features

Given the spectral and complexity differences between WM and GM, we tested if matter type could be decoded from the extracted signal features using an ensemble of k-nearest neighbor (KNN) classifiers. The model was trained on one (exponent) or five features (exponent, delta and alpha central frequency, HFD, LZC). The subspace ensemble consisted of 30 KNN learners where each ensemble was trained on a random subset of predictors with a subspace dimension of three features max per learner (s = 3). Model performance was evaluated using a 5-fold cross validation. To account for different numbers of WM and GM contacts, we up-sampled the WM by duplicating a randomized subset of contacts, a standard approach for unbalanced data (Sun, Wong and Kamel, 2009). Using the exponent as the only feature, our model achieved an AUC = 0.76 (permutation test, n = 1000, p<0.001). Including delta central frequency, alpha central frequency, HFD, and LZC as well, we achieved an AUC = 0.92 (permutation test, n = 1000, p<0.001), indicating that these signal features provide non-redundant information and lead to a more accurate separation between signals in WM and GM (Figure 2F).

### Complexity of WM signals is related to their structural features

For a subset of participants (n=9), we tested whether the complexity of the signals recorded from WM locations related to structural integrity of the surrounding WM, as indexed by FA. We found a weak positive correlation between FA and HFD (rho = 0.12, p=0.023; Figure 3A), and just a trend for FA-LZC (rho = 0.09, p=0.098). To further assess the robustness of the HFD-FA relationship across our sample, we fitted a mixed-effects model with HFD as the dependent variable, FA as the independent variable, and participants as random effects. We found that higher FA values predict higher HFD scores (β = 0.108, p=0.0075).

**Figure 3.**
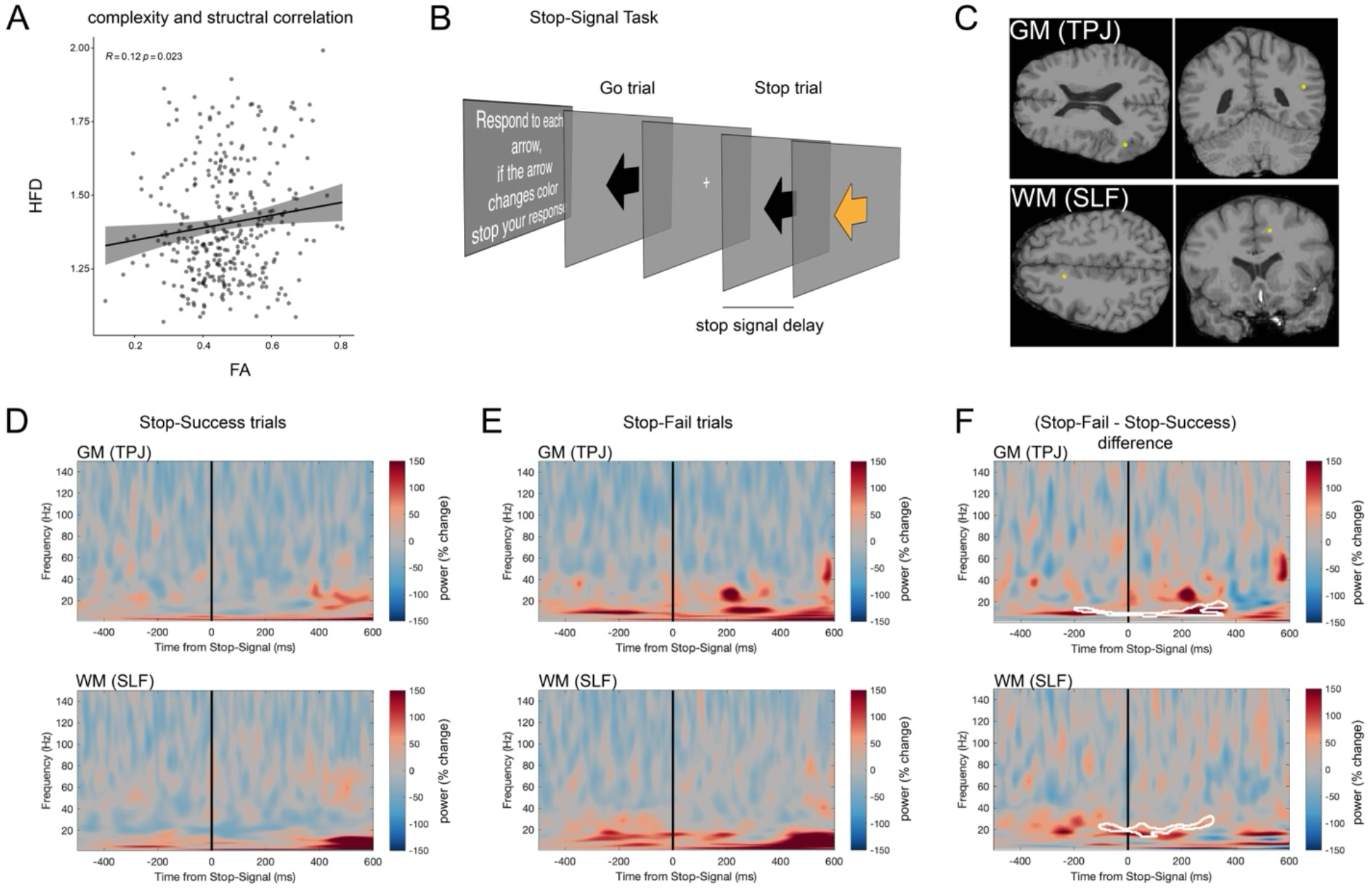
Structural and functional WM effects. Panel A: weak positive correlation between FA and HFD, with each WM electrode contact displayed as a circle with the correlation fit line overlaid. Panel B: schematic of the stop-signal task, with go and stop trials. Panel C: example of an electrode location in GM (temporo-parietal junction, TPJ) and in WM (Superior Longitudinal Fasciculus, SLF) displaying task-related modulations (patient 5). Panel D: Stop-Success average power values (percent change from baseline) across the time-frequency space (500 before to 600 ms after the stop signal, frequencies from 2 to 150 Hz) for the GM and WM locations shown in the previous panel, displayed on top and bottom respectively. Panel E: Same as D for Stop-Fail. Panel F: same as previous panels displaying the difference in power values between Stop-Fail and Stop-Success trials, with regions of significant statistical difference based on permutation testing outlined in white (p-corrected<0.05).

### Control Analyses with a different re-referencing scheme

To confirm our results did not depend on a specific signal referencing scheme, we repeated all analyses after employing an alternative approach to re-reference all signals. The average of all WM signals was used to re-reference WM signals, and the average of all GM signals to re-reference GM signals. In brief, the spectral and complexity values were very similar to those obtained with the common average reference, and all the findings were replicated. WM displayed a lower overall power (WM offset = 2.93 ± 0.921; GM offset = 3.88 ± 0.843; β = −0.94, p<0.001), a shallower 1/f shape (WM exponent = 2.45 ± 0.392; GM exponent = 2.78 ± 0.365; β = −0.31, p<0.001), and higher central frequencies for periodic components in the delta and alpha range (WM delta range central frequency = 2.39 ± 1.14 Hz; GM delta range central frequency = 1.99 ± 1.10 Hz; β = 0.42, p<0.001; WM alpha range central frequency = 11.0 ± 1.59 Hz; GM alpha range central frequency = 10.7 ± 1.53 Hz; β = 0.35, p<0.001). The only difference with respect to previous results was that the effect also reached significance in the theta range (WM theta range central frequency = 6.39 ± 1.16 Hz, GM = 6.36 ± 1.20 Hz, β = 0.14, p=0.0128). WM signals were more complex (HFD WM: 1.39 ± 0.189; HFD GM: 1.29 ± 0.1691; β = 0.112, p<0.001; LZC WM: 0.467 ± 0.279; LZC GM: 0.348 ± 0.228; β = 0.127, p<0.001) and complexity correlated negatively with the exponent values (HFD-exponent rho = −0.60, p<0.001; LZC-exponent: rho = −0.19, p<0.001). When considering complexity and FA covariations, higher FA predicted higher HFD values, but not LZC, like in the main results (FA-HFD: rho = 0.14, p=0.013; HFD by FA mixed effect model: β = 0.134, p=0.0024; FA-LZC: rho = 0.08, p=0.16).

### WM signals show functional activity

Finally, for a subset of participants we performed a time-frequency analysis of signals recorded from WM locations during the stop-signal task (Figure 3B). We compared power values associated with Stop-Success and Stop-Fail (permutation testing with cluster-based multiple comparison correction, corrected-p<0.05) to evaluate if we could detect behaviorally-relevant differences in activity recorded from WM locations. In 6 out of 9 participants at least one WM location displayed a difference between Stop-Success and Stop-Fail trials, indicating that functional, task-related signals can be detected in WM locations. The proportion of WM electrodes with signals displaying task-related modulations ranged between 0-39% across patients, resembling the proportion found when performing the same analysis on GM signals (0-29%). This further suggests that WM functional signals are not ubiquitous to all WM but depend on the circuit engaged by the task and by the proportions of electrodes sampling that circuit for any given participant, just like GM effects. The task-modulations for both WM and GM occurred in the broadband gamma range (>40 Hz) or in the beta range. The WM locations displaying task modulations spanned several tracts (Table 1): the Uncinate Fasciculus (UF), the Inferior Fronto-Occipital Fasciculus (IFOF), the Inferior Longitudinal Fasciculus (ILF), the Arcuate Fasciculus (AF), the Frontal Aslant Tract (FAT), the Superior Longitudinal Fasciculus (SLF), Forceps Minor (FM), and Optic Radiation (OR). Specifically, Stop-Success was associated with increased activity in the UF, IFOF, and ILF. Failures to stop were associated with activity in the FM, FAT, and late activity in OR. Both the AF and SLF displayed variability in the direction of the effects, with increased activity for both Stop-Success and Stop-Fail. Given the presence of different subdivisions of the SLF, with three different major bundles, it is possible that the variability in activity reflects the different functional subdivisions of the different bundles. In Figure 3C-F we showcase two distant electrode contacts in one participant (patient 5; Figure 3C) – one in GM (temporo-parietal junction, TPJ) and the other in WM (SLF) – and we display their average activity during Stop-Success (Figure 3D) and Stop-Fail trials (Figure 3E). Both locations are characterized by increased and sustained beta-band activity for Stop-Fail trials, starting before the stop signal, with a very similar spectro-temporal profile (Figure 3F). The similarity observed in this pairs of distant recording sites may be explained by the fact that TPJ projects to frontal cortex through the SLF.

**Table 1.**
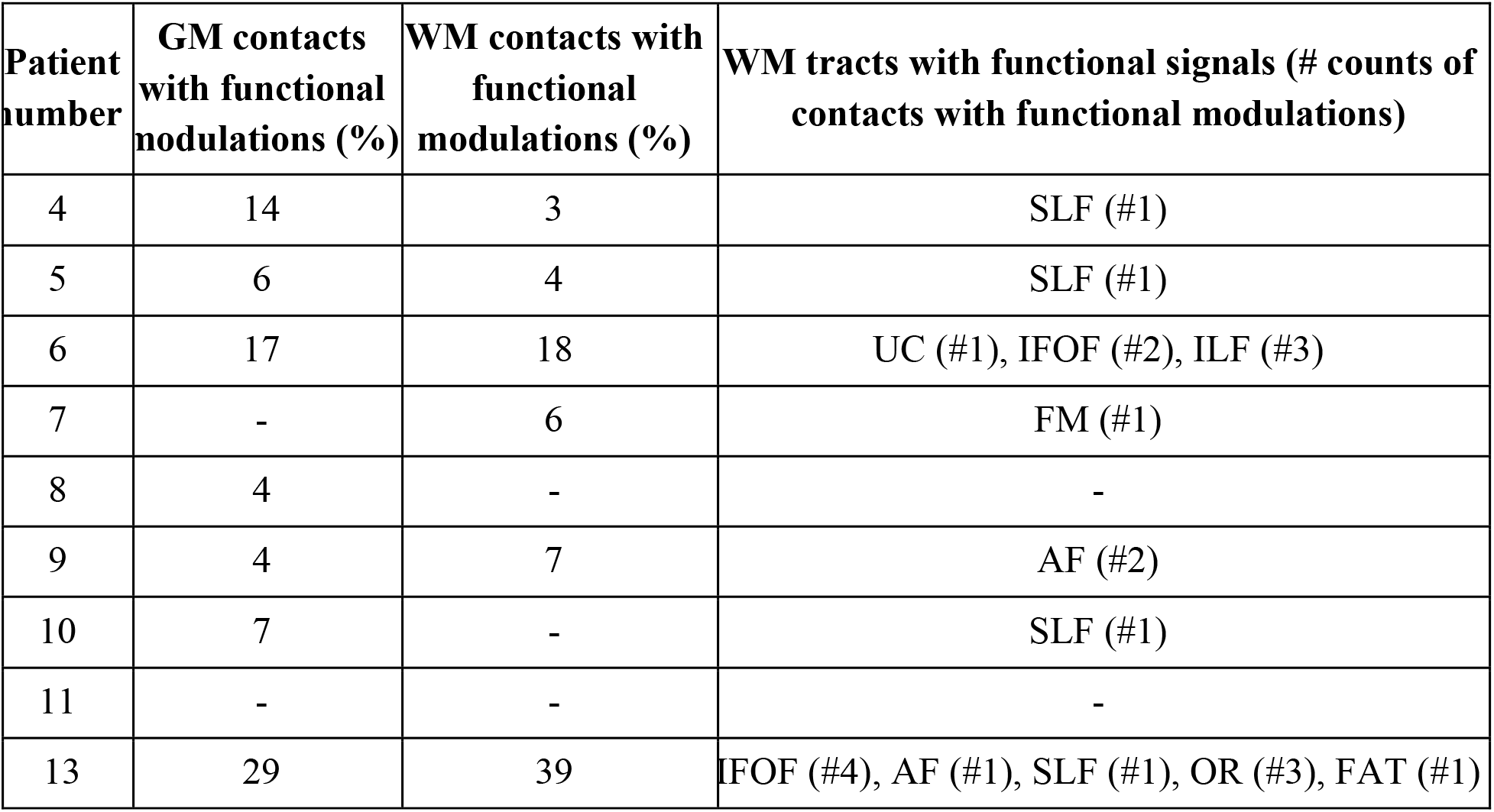
Functional signals (difference between Stop-Success and Stop-Fail trials in the stop-signal task) in GM and WM locations for each participant. Columns report the proportion of electrodes with functional differences with respect to the total electrodes analyzed for GM and WM, and the putative tracts associated with the WM locations (number of electrodes for each tract reported in parenthesis).

| **Patient number** | **GM contacts with functional modulations (%)** | **WM contacts with functional modulations (%)** | **WM tracts with functional signals (# counts of contacts with functional modulations)** |
| --- | --- | --- | --- |
| 4 | 14 | 3 | SLF (#1) |
| 5 | 6 | 4 | SLF (#1) |
| 6 | 17 | 18 | UC (#1), IFOF (#2), ILF (#3) |
| 7 | - | 6 | FM (#1) |
| 8 | 4 | - | - |
| 9 | 4 | 7 | AF (#2) |
| 10 | 7 | - | SLF (#1) |
| 11 | - | - | - |
| 13 | 29 | 39 | IFOF (#4), AF (#1), SLF (#1), OR (#3), FAT (#1) |

## Discussion

In line with our hypothesis, our results indicate that signals recorded from WM have distinct spectral features from those recorded from GM, and that this difference cannot be simply attributed to an attenuation of the signal recorded from WM. Indeed, while WM signals were lower in power than GM (lower offset derived from the spectral parametrization model), they displayed additional differences in both aperiodic and periodic activity, and they also exhibited differences in signal complexity metrics which correlated with WM structural characteristics. Finally, we found evidence of functional information in signals recorded from WM, in the form of behavior-related modulations, potentially indicating a read-out of network-wide activity through WM tracts.

When considering the spectral components, WM signals displayed a different distribution of power across frequencies compared to GM, in the form of a shallower 1/f slope (lower exponent values in 1/f^*x*^). Relative to GM, this translates to reduced low-frequency power and elevated high high-frequency power, indicating more asynchronous activity in the local field potential (Donoghue *et al*., 2020). Investigations of the 1/f slope are becoming increasingly popular in clinical research on neurological and psychiatric diseases (Donoghue, 2025; Hacker *et al*., 2025). Relevant to our work, one study reported longitudinal spectral slope changes as a biomarker for treatment efficacy in treatment-resistant depression, measured through chronic deep brain stimulation (DBS) device recordings from the subcallosal cingulate (Veerakumar *et al*., 2019). The subcallosal cingulate is a WM location at the intersection of four different WM tracts, the UF, FM, cingulum and fronto-striatal fiber bundles (Tsolaki *et al*., 2021), and DBS is believed to mediate the antidepressant response through the engagement of said tracts. Thus, existing evidence regarding spectral slope measured at WM locations is strongly suggestive that WM signals contain clinically relevant information reflecting WM tract activity.

Periodic components also displayed differences between WM and GM, with a shift in the frequency of delta and alpha-range oscillations toward higher values, thus reflecting relatively faster oscillations in WM. While the origin and circuit-level interpretation of these changes requires further investigations and biophysical modeling to be understood, previous evidence demonstrated that functional connectivity, measured intracranially using WM signals across various canonical oscillations ranges, was strongly correlated with fMRI-derived functional connectivity (Huang *et al*., 2023). Investigations linking neuronal firing rate changes to nearby oscillatory activity (Jourahmad *et al*., 2026) could be applied to test if WM periodic signals from a given tract show any reliable relationship with neuronal activity measured in the interconnected GM regions.

We further quantified signal complexity in terms of unpredictability and irregularity of the time-series using HFD and LZC (Lau *et al*., 2022), finding that WM signals were more complex than GM ones. While this could appear counterintuitive at first, it is expected based on the aperiodic activity results. Indeed, complexity and spectral slope have been shown to be related to each other using simulations and recordings from multiple species (Medel *et al*., 2023). We replicate this finding here, with shallower 1/f slope values corresponding to increased complexity scores, a relation reliably observed at the single subject level. Intuitively, the more asynchronous activity leading to the shallower spectral slope would also correspond to less predictable signals. Finally, when using the 1/f spectral slope values, a decoder was able to classify WM from GM with 76% accuracy. When periodic and complexity features were also included, the ability to classify WM versus GM signals improved to above 90%, indicating that while spectral and complexity features are related to each other, they still provide non-redundant information and lead to an improved classification accuracy when considered jointly.

To evaluate whether the signal complexity reflected sparsely sampled neural activity traveling along WM tracts, we tested whether complexity related to the structural features of the WM locations from which we recorded, by means of FA. We found that, at a group level, HFD was positively correlated with FA, indicating that WM locations associated with highly organized, structurally intact, and well-defined white matter tracts, displayed electrophysiological signals with higher complexity. This result resembles similar previous findings obtained with magnetoencephalography, where an increase in complexity positively correlating with FA was observed in a subset of WM tracts using source localization (Fernández *et al*., 2011). Overall, this suggests that the temporal patterns of WM signals are influenced by WM structural features, and might be a novel avenue to investigate and detect WM degradation in diseases characterized by demyelination.

After broadly characterizing WM and GM signal features using resting state recordings, we looked into their functional information content by considering recordings during the performance of a task. Functional neuroimaging studies reported circuit-specific task-related activity in WM tracts more than 20 years ago (Tettamanti *et al*., 2002), and the evidence of task-locked BOLD signals within WM tracts is vast (Mazerolle *et al*., 2010; Tae *et al*., 2014; Ding *et al*., 2018; Li *et al*., 2019; Schilling *et al*., 2023). However, the different hemodynamic characteristics of WM BOLD responses (Fraser *et al*., 2012; Courtemanche *et al*., 2018) have been a source of confusion and these effects, while overwhelmingly present, are typically underreported in the neuroimaging literature (Tettamanti *et al*., 2002; Yarkoni *et al*., 2009; Gawryluk, Mazerolle and D’Arcy, 2014; Spencer *et al*., 2025). A similar situation might be occurring in electrophysiological studies using sEEGs. To our knowledge, only one previous sEEG study explicitly studied WM signals for their functional significance, showing that the inclusion of WM signals in conjunction with GM signals was able to improve the decoding of movement (Li *et al*., 2021). In our own experience with sEEG recordings, we often noticed task-related activity in electrodes later recognized to be in WM, motivating the present study. Here, we found task-related modulations across several WM locations, with similar proportions of effects to those obtained across GM locations. During the successful inhibition of a motor response in a stop-signal task, we found increased activity in the ILF, UF and IFOF. These tracts support the propagation perceptual information and attentional resources between the occipital (ILF and IFOF) and temporal (UF) lobe and the frontal cortex (Bullock *et al*., 2022). In the frontal cortex, targets include the inferior frontal gyrus and orbitofrontal cortex (Giampiccolo, Herbet and Duffau, 2025), two key frontal regions for stopping (Aron, Robbins and Poldrack, 2004, 2014; Hampshire *et al*., 2010; Bartoli, Aron and Tandon, 2018; Balasubramani, Pesce and Hayden, 2020). When inhibition failed and the motor response could not be interrupted, we observed activity in the FM, a frontal interhemispheric tract which might underlie a bilateral engagement of prefrontal activity, opposing the lateralized recruitment of right prefrontal cortex necessary for successful stopping (Aron, Robbins and Poldrack, 2004), and in the FAT, a tract connecting presupplementary areas to the inferior frontal gyrus, and believed to play a key role in motor preparation and its control (Budisavljevic *et al*., 2017; Landers *et al*., 2022). The engagement of these tracts during Stop-Fail trials might reflect the uncontrolled release of the prepotent motor plan. The SLF, connecting parietal and frontal cortex, displayed variable signatures, likely due to its subdivisions and their different functional roles (Janelle *et al*., 2022), in line with recent findings on parietal cortex’s role in stopping and its functional connectivity profile with different prefrontal regions (Osada *et al*., 2019; Kang *et al*., 2025; Kang, 2026). Future studies carefully examining functional activity related to WM tracts at the group-level will be necessary to fully understand the extent of these promising indications of functional signals in WM.

A few limitations should be considered. The spectral and complexity effects reported were obtained considering WM and GM differences overall, regardless of their precise anatomical origin. It is likely that regional, tract-specific effects exist, and that WM signal properties are further influenced by the location of the electrode contact with respect to the local microscopic and macroscopic anatomy, for example their position with respect to axonal orientation. Future studies could systemically compare signal features obtained from WM while carefully considering their anatomical properties. Complexity metrics were evaluated only during resting state, and more research is needed to understand how spectral and complexity relationships evolve during state perturbations, like active behavior. Additionally, the functional effects were evaluated at the single-subject level due to the sample size and the sparse sampling of different WM tracts across different patients. Group-level analysis of WM functional signals will require recruiting participants according to their WM tracts coverage to ensure sufficient comparability of WM locations and their signals, and with ad-hoc hypotheses about their functional role (Giampiccolo, Herbet and Duffau, 2025). Finally, signal features might be affected by the re-referencing scheme employed. Mercier et al. carefully investigated WM versus GM signals across different re-referencing schemes, finding that WM locations displayed unique activity with respect to GM even when using the most conservative approaches (Mercier *et al*., 2017). In our control analysis, our results were robust and virtually identical across the two different re-referencing schemes tested.

Crucially, our results indicate that a common assumption, that contacts in WM signals can be used as a neutral recording reference in sEEG recordings, might need to be re-evaluated. We recently demonstrated how relying on contacts in WM for the recording reference can bias estimates of effective connectivity obtained through cerebro-cerebral evoked potentials (Huang *et al*., 2025), and that a modified common average re-referencing scheme (Huang *et al*., 2024) is necessary to remove the bias and obtain reliable results. Using an extracranial, subgaleal reference might represent a valid alternative to obtain a neutral recording reference (Devara *et al*., 2026).

In summary, our results indicate that electrophysiological signals recorded from WM locations display distinct features with respect to GM and they carry complex, task-relevant information. Thus, they should not be discarded during analysis, as they might offer a window into the information travelling along WM tracts (Mercier *et al*., 2017; Mercier, 2026; Revell *et al*., 2026). WM tracts compose over 40% of brain volume, and their integrity supports the coordination and balance of activity between GM regions and nuclei. Future investigations into the functional and clinical relevance of WM signals could become critical for advancing our knowledge of network-level disorders, including several psychiatric and neurological conditions (Giampiccolo *et al*., 2026; Mercier, 2026; Revell *et al*., 2026).

## Data availability statement

Data will be available upon reasonable request.

## Acknowledgments and Conflict of Interest disclosure

We thank the patients for their participation and the hospital staff for their assistance.

S.A.S has consulting agreements with Boston Scientific, NeuroPace, Koh Young, Zimmer Biomet, Abbott, and is co-founder of Motif Neurotech.

