## Supplementary Material for "Electrophysiological features of signals recorded from white matter"

| Patient number | Age | Sex | Cortical GM contacts | WM contacts | Diffusion MRI | Stop-Signal task |
|---|---|---|---|---|---|---|
| 1 | 24 | M | 99 | 54 | y | - |
| 2 | 52 | F | 50 | 30 | - | - |
| 3 | 23 | M | 72 | 8 | - | - |
| 4 | 44 | F | 65 | 37 | y | y |
| 5 | 39 | F | 34 | 26 | - | y |
| 6 | 46 | F | 69 | 37 | - | y |
| 7 | 42 | M | 30 | 30 | - | y |
| 8 | 25 | M | 96 | 35 | y | y |
| 9 | 40 | F | 55 | 28 | y | y |
| 10 | 55 | F | 32 | 21 | y | y |
| 11 | 30 | M | 24 | 20 | - | y |
| 12 | 63 | M | 72 | 5 | - | - |
| 13 | 44 | F | 39 | 27 | y | y |
| 14 | 24 | M | 70 | 46 | y | - |
| 15 | 32 | F | 61 | 56 | y | - |
| 16 | 52 | M | 77 | 30 | - | - |
| 17 | 20 | M | 76 | 33 | y | - |
| 18 | 25 | F | 60 | 42 | - | - |
| 19 | 41 | F | 18 | 53 | - | - |
| **Total** | | | **1099** | **618** | | |

*Table S1. Demographic and electrode information for each participant. The subset of participants with diffusion MRI information and those who participated in the stop-signal task are indicated in the last two columns.*
